# Mitochondrial heteroplasmy is responsible for Atovaquone drug resistance in *Plasmodium falciparum*

**DOI:** 10.1101/232033

**Authors:** Sasha Siegel, Andrea Rivero, Swamy R. Adapa, ChengQi Wang, Roman Manetsch, Rays H.Y. Jiang, Dennis E. Kyle

## Abstract

Malaria is the most significant parasitic disease affecting humans, with 212 million cases and 429,000 deaths in 2 015^1^, and resistance to existing drugs endangers the global malaria elimination campaign. Atovaquone (ATO) is a safe and potent antimalarial drug that acts on cytochrome *b* (cyt. b) of the mitochondrial electron transport chain (mtETC) in *Plasmodium falciparum,* yet treatment failures result in resistance-conferring SNPs in cyt. b. Herein we report that rather than the expected *de novo* selection of resistance, previously unknown mitochondrial diversity is the genetic mechanism responsible for resistance to ATO, and potentially other cyt. *b* targeted drugs. We found that *P. falciparum* harbors cryptic cyt. *b.* Y268S alleles in the multicopy (∼22 copies) mitochondrial genome prior to drug treatment, a phenomenon known as mitochondrial heteroplasmy. Parasites with cryptic Y268S alleles readily evolve into highly resistant parasites with >95% Y268S copies under in vitro ATO selection. Further we uncovered high mitochondrial diversity in a global collection of 1279 genomes in which heteroplasmic polymorphisms were >3-fold more prevalent than homoplasmic SNPs. Moreover, significantly higher mitochondrial genome copy number was found in Asia (e.g., Cambodia) versus Africa (e.g., Ghana). Similarly, ATO drug selections *in vitro* induced >3-fold mitochondrial copy number increases in ATO resistant lines. Hidden mitochondrial diversity is a previously unknown mechanism of antimalarial drug resistance and characterization of mitochondrial heteroplasmy will be of paramount importance in combatting resistance to antimalarials targeting the electron transport chain.

---

Atovaquone (ATO), a napthoquinone, and the pyridones^2-4^ acridones^5^, Acridinediones^6-8^, tetrahydroacridines^6^, and the 4(1H)-quinolones^9-12^ potently inhibit the cytochrome *bc1* complex of the mitochondrial electron transport chain (mtETC) within the mitochondria, with disruption of pyrimidine biosynthesis and collapse of mitochondrial membrane potential leading to parasite death^13^. The first ATO treatment failures were observed in the Phase II clinical trials between 1991-1994 in Thailand^14^. These studies demonstrated that ATO monotherapy resulted in clinical treatment failures and subsequent recrudescence of infection, prompting the use of ATO in combination with proguanil (AP) for malaria prophylaxis and treatment. The clinical experience with ATO and numerous *in vitro* drug selection studies led to a hypothesis that ATO resistance arises readily *de* novo following treatment^15-17^, however *in vitro* data are incongruent with clinical data of a single Y268S/N/C SNP as the genetic mechanism in two important aspects. First, clinical treatment failures are most commonly linked to an amino acid substitution at position Y268 in cyt. b^18^ but *in vitro* drug pressure selects for a variety of different mutations except Y268: M133I, M133V, P275T, K272R, G280D, L283I, V284K, L144S and F267V^18-20^ (Extended Data Table 1). Second, drug susceptibility studies with ATO resistant Y268S mutants demonstrate a broad range of potency rather than a dichotomous response that would be expected from a single SNP. By using paired parasites collected upon patient admission and subsequent treatment failure from Phase II trials of ATO (Extended Data Table 2), we discovered three distinct *in vitro* ATO resistance phenotypes (Table 1). A recrudescent isolate (TM90-C6B) initially typed as Y268 (WT) exhibited low level ATO resistance, other isolates with Y268S from recrudescent isolates (e.g., TM90-C2B) demonstrated moderate resistance to ATO and myxothiazol, and two isolates from ATO-pyrimethamine treatment failures showed extreme resistance (TM92-C1086 and TM92-C1088) (Table 1). Surprisingly, the extreme ATO resistance phenotype produced greatly reduced susceptibility to a broad range of mtETC inhibitors (Table 1), even though the parasites expressed the common Y268S/N SNPs and no apparent resistance-associated SNPs in candidate mtETC encoding genes (Extended Data Table 3). These data demonstrate that more than a single cyt. *b* SNP mediates the different ATO resistance phenotypes.

**Table 1.** Drug susceptibility (IC_50_s, nM) of *Plasmodium falciparum* isolates and clones to mitochondrial electron chain inhibitors reveal three atovaquone (ATO) resistance phenotypes^*^

|  |  |  | Complex III |  |  |  |  |  |  |  |  |
| --- | --- | --- | --- | --- | --- | --- | --- | --- | --- | --- | --- |
|  | NDH2 enzyme | DHODH enzyme | Qo site (cyt *b*) |  | Qi site (cyt *b*) |  |  | unknown |  |  |  |
|  | HDQ | DSM-1 | ATO | MYX | ANT | ELQ 300 | GSK121 | ICI56,780 | P4Q-391 | Cyt. *b* genotype | Admission/<br>Recrudescent |
| W2 | 55.0 | 52.4 | 0.41 | 10.8 | 165.5 | 2.72 | 3.39 | 0.03 | 11.1 | WT | -- |
| TM90-C6B | ND | 24.5 | 109 | 7.45 | 125 | 7.67 | 36.4 | 0.50 | 33.0 | WT | R |
| TM90-C2A | 14.8 | 79.4 | 3.09 | 154 | 72.0 | 0.69 | 20.2 | 0.04 | 1.92 | WT | A |
| TM90-C2B | 146 | 57.4 | 5290 | 428 | 152 | 4.61 | 77.5 | 14.3 | 55.5 | Y268S | R |
| TM90-C40B2 | 830 | 47.2 | 1.53 | 45.1 | 72.9 | 4.02 | 7.84 | 0.04 | 22.1 | WT | A |
| TM90-C50B5 | ND | ND | 3940 | 3010 | ND | 6.45 | 122 | 13.4 | 32.4 | Y268S | R |
| TM92-C1086 | 180 | 770 | 31400 | 4160 | >18200 | 16300 | 2470 | 882 | 14160 | Y268S | R |
| TM90-C1088 | 252 | 540 | 29100 | 3990 | >18200 | 21000 | 3470 | 219 | >20200 | Y268S | R |

\*Atovaquone (ATO) Resistance phenotypes: low (light grey), moderate (grey), and extreme (dark grey); Drugs: HDQ – 1-hydroxy-2-dodecyl-4-(1H)-quinolone; MYX – myxothiazol; DSM-1 – triazolopyrimidine; ANT – antimycin A; GSK121 – GSK932121A, 4(1H)-pyridone; ICI56,780 – phenoxyethoxy-4(1H)-quinolone; P4Q-391 – 4(1H)-quinolone

The mitochondria of asexual erythrocytic stage *P. falciparum* exist as a single organelle with a linear, non-recombining, tandemly-repeated 6-kb genome with approximately 22 copies present in each parasite, and is remarkably well-conserved compared to the nuclear and apicoplast genomes^21^. These data suggest there are inherent mechanisms in place to conserve the mitochondrial genome that are exclusive to the mitochondria and its three encoded genes: cyt. *b,* cytochrome c oxidase subunit I (*cox I*), and cytochrome c oxidase subunit III (*cox III*)^21^. The existence of multiple mtDNA copies creates the potential for mitochondrial allelic heterogeneity within a single parasite as well as the mtDNA pool at the population level, a phenomenon known as mitochondrial heteroplasmy^22^. Therefore, we hypothesized mitochondrial heteroplasmy as a genetic mechanism for ATO resistance and broader mitochondrial diversity.

If heteroplasmy is a source of mitochondrial diversity, parasites of different genetic backgrounds could possess variant SNPs at very low levels not detected by conventional Sanger sequence genotyping methods. To assess presence of heteroplasmic SNPs that confer ATO resistance, we used a pyrosequencing assay for quantitative analysis of cyt. *b^23^* to quantify the Y268S allele (Figure 1A) and applied it to the earliest cryopreserved *P. falciparum* isolates from the Phase II studies (Extended Data Table 1). These data indicate the presence of cryptic mutant alleles present in admission parasite isolates (e.g., TM90-C2A and TM90-C40B2), with ∼1-2% mutant Y268S frequency detected (Figure 1B). Similarly, ATO resistant isolates (e.g.,TM90-C2B and TM90-C50B5) from clinical failures possessed extremely high copies of Y268S. The TM90-C6B isolate had low-level Y268S presence (Figure 1B); interestingly this isolate was from a clinical failure initially identified as WT Y268 cyt. *b* (Extended Data Table 1). In contrast, no Y268S heteroplasmy was found in *P. falciparum* clones unable to develop ATO resistance (e.g., D6) ^24^ or found to develop alternative cytochrome *b* mutations (e.g., W2; Extended Data Table 2), ^18,20^ confirming that the observed Y268S allele is only present at low levels in some parasite mitochondrial haplotypes.

**Figure 1.**
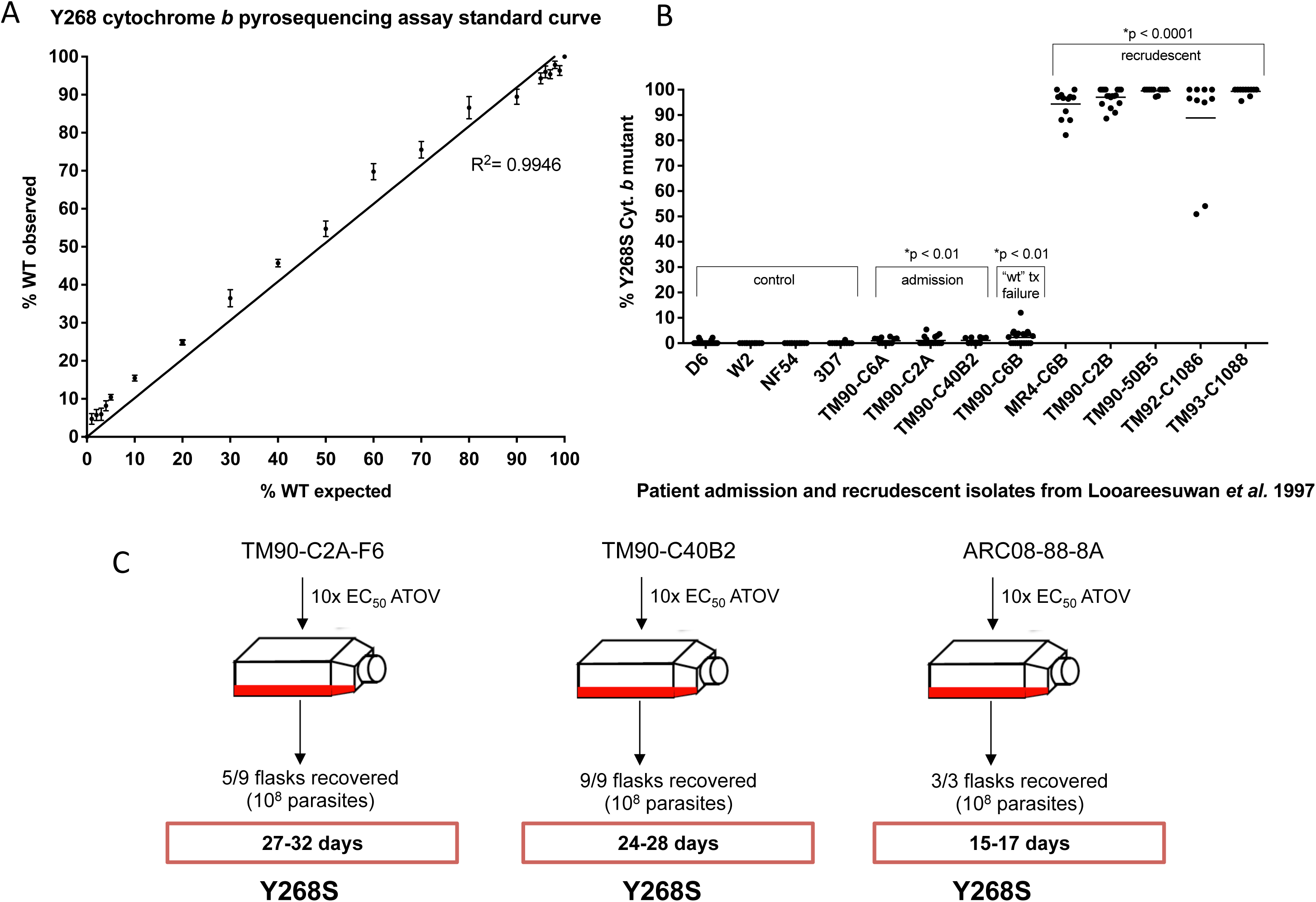
Pyrosequencing assessment of Y268S allele in patient isolates and drug selection strategy. A. Y268S cytochrome *b* pyrosequencing assay standard curve reliably detects frequencies of wild-type and mutant alleles. Using wt and mutant gDNA mixtures from 0% to 100% wt, the standard curve shows the correlation between the percentage of wt DNA expected in each mixture and the pyrosequencing assay’s ability to detect that percentage, with a correlation coefficient of R^2^=0.9946. B. Pyrosequencing analysis of the Y268S allele in admission and recrudescent isolates from Phase II studies in Thailand. Pyrosequencing analysis of control lab strains that lack the Y268S allele indicate that there is a very low false positive detection rate using this assay. Admission isolates show low level Y268S heteroplasmy, as well as “wt” treatment failure TM90-C6B, and recrudescent parasites show high levels of the Y268S allele. C. *In vitro* drug selections with clinical isolates harboring cryptic Y268S alleles. Admission isolates from the Phase II studies of ATO in Thailand (TM90-C2A and TM90-C40B2) and isolate from Thailand in 2008 (ARC08-22-4G) all had low level Y268S heteroplasmy, and rapidly developed majority Y268S mtDNA after exposure to 10xEC50 ATO, recovering after 15-32 days

To independently investigate the novel parasite heteroplasmy genetics, we next studied the same set of isolates by Illumina sequencing with very deep coverage (10,000-30,000x coverage of mtDNA) and determined the Y268S allele frequency (AF) (Figure 2A). These data confirmed the presence of low-level Y268S heteroplasmy in *P. falciparum* isolates prior to treatment and in a 2008 isolate from Cambodia (PL08-025). The deep sequencing data for recrudescent ATO resistant isolates were consistent with the pyrosequencing analysis, demonstrating very high, yet not purifying levels of Y268S selection in the mtDNA. Interestingly, ATO resistant TM93-C1090 possessed low-level Y268S although the majority of mitochondrial alleles were Y268N. The corresponding admission isolate for this parasite (TM93-C0151) only showed low-level Y268S. It is plausible that this isolate had *de novo* selection of Y268N or possessed undetectable Y268N (< 0.5% AF) pre-treatment.

**Figure 2.**
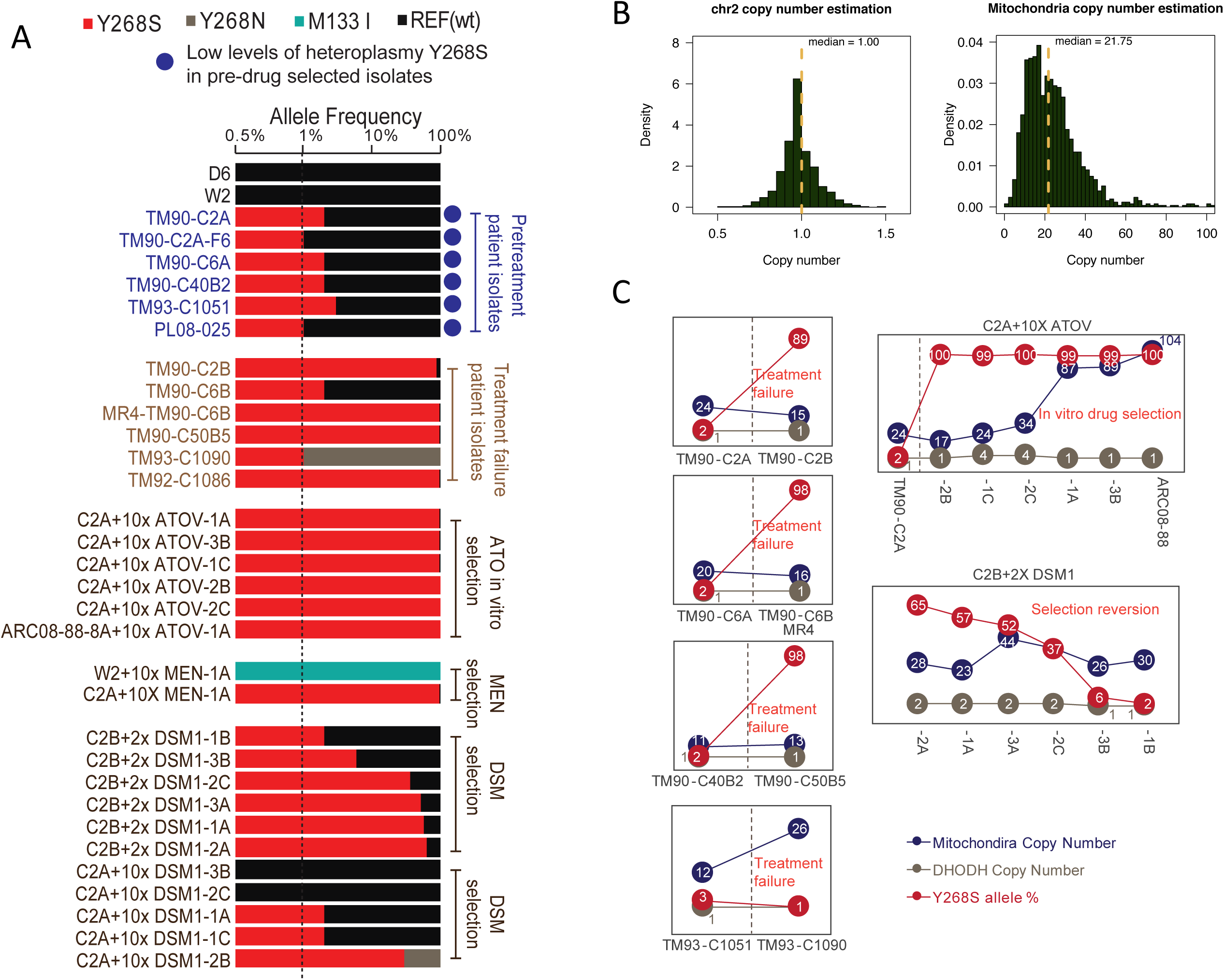
Using mitochondrial deep sequencing to identify low frequency heteroplasmy sites in patient isolates and from in vitro drug selections. A. Deep sequencing shows heteroplasmy/homoplasmy status of patient admission/recrudescent isolates and *in vitro* drug selected parasites at known ATO resistance alleles. Parasites were taken from earliest cryopreserved isolates and mtDNA was sequenced at > 10,000x coverage to preserve respective phenotypes/genotypes. B. Histogram plot shows the distribution of estimated copy number of nuclear and mitochondria genomes (yellow line indicates the median value). For 1279 *P. falciparum genome* samples, we extracted median coverage (DP) for each reported sites in chromosome 2 (*N*, median value 1.00) and mitochondria genome (*M*, median value 21.75). C. Heteroplasmy and copy number from clinical isolates and *in vitro* drug selection studies. Both Y268S heteroplasmy and copy number variations were found in *in vitro* drug selection studies

Overall, these findings sharply contrast with previous reports of *de novo* Y268S mutant selection in patients treated with ATO, since Y268S heteroplasmy demonstrates mutant alleles already were present in admission isolates prior to ATO exposure, but it required very high coverage, PCR-free Illumina sequencing of the mtDNA to uncover these low level heteroplasmic alleles. Therefore, we next aimed to experimentally demonstrate the potential for ATO resistance to result from low-level Y268S heteroplasmy *in vitro*, which to our knowledge, has never been successfully generated in prior studies. We selected for resistance to ATO in *P. falciparum* clones isolated from patients prior to ATO treatment (TM90-C2A and TM90-C40B2) and a 2008 clone from Cambodia (ARC08-88-8A; Figure 1C); each of these expressed low level Y268S heteroplasmy in pyrosequencing and deep sequencing analysis. Parasites were seeded at 10^8^ per flask and exposed to 10x EC_50_ concentrations of ATO, and within 15-27 days all generated resistance with cyt. *b* Y268S mutations (Figures 1C and 2A). We then investigated if the Y268S mutation could be induced by another cyt. *b* acting drug, menoctone (MEN), a 2-hydroxy-3-(8-cyclohexyloctyl)-1,-4-naphthoquinone thought to have the same mechanism of action as ATO^25^. MEN at 10x EC_50_ (1.5 μM) selected for Y268S in TM90-C2A, but W2 developed the common M133I mutation associated with ATO exposure in other lab strains (Extended Data Figure 3) ^18,20,26^ High coverage deep-sequencing results confirmed 99% Y268S and M133I frequencies in these drug selected mutants (Figure 2A). The MEN resistance phenotype of C2A+10xMEN-1A displayed high-grade ATO resistance with an EC50 of 54 μM, similar to that of TM92-C1086. Conversely, W2+10xMEN-1A expressed moderate resistance (EC_50_ = 66 nM), presumably reflective of fitness differences between cyt. *b* Y268S and M133I. These data are consistent with other M133I selected mutants^18^ (Extended Data Table 2). Obtaining the clinically relevant Y268S mutation in an *in vitro* setting using multiple drugs shows that pre-existing, low-level heteroplasmic SNPs are responsible for the development of resistance in these parasites.

*P. falciparum* dihydroorotate dehydrogenase (DHODH) is the only essential enzyme in the mtETC that is required to synthesize pyrimidine precursors, and we hypothesized that parasites with resistance to both DHOD and cyt. *b* inhibitors could recapitulate the extreme resistance phenotype. DSM1, a potent inhibitor of DHODH, induces mutations in DHODH as well as increased copies of the DHODH gene^19,27^. Unsurprisingly, 10x EC_50_ DSM1 pressure in the ATO sensitive TM90-C2A background readily generated DSM1 resistant parasites with copy number amplification in DHODH (Extended Data Table 4). However, there were two unusual selections in this background, with C2A+10xDSM-2B being a mixed genotype of 30% Y268S and 8% Y268N, a tri-allelic heteroplasmic parasite being selected by DSM1 alone (Figure 2A). In addition, this parasite was a partial R265G mutant in DHODH (34%); this mutation was only observed in one other DSM1 selected parasite from TM90-C2A (C2A+10xDSM1-3B), that was predominantly an R265G mutant (97%). This raises the possibility that DSM1 exposure can induce increased heteroplasmy in cyt. b., although in an unstable and inefficient manner.

*In vitro* DSM1 drug resistance selection studies in an ATO resistant background yielded intriguing results on the role of heteroplasmy and mitochondrial copy number. DSM1 drug selections with ATO resistant TM90-C2B were not successful at 10x EC_50_ (1.5 uM) concentrations, with three failed attempts (Extended Data Figure 4). Using lower DSM1 concentrations (2x EC_50_ = 300 nM) made resistance development possible. Deep-sequencing of C2B+10xDSM1 selected lines (Figure 2A) revealed that parasites thawed from earliest cryopreserved line and expanded for sequencing (45 days of total DSM1 exposure) largely lost their initial resistance to ATO during DSM1 exposure, with parasites gradually exchanging their Y268S allele in favor of copy number amplifications in DHODH, implying that these resistance mechanisms are antagonistic at low concentrations of DSM1, but completely incompatible at high concentrations. The initial ability to survive both DSM1 and atovaquone pressure could consist of immediate reactions to respirational stress, where DSM1 induces DHODH amplifications quickly and proportionately to DSM1 exposure, and Y268S is cryptically present and able to increase allele numbers at any time. Both of these mechanisms are attractive from the metabolic standpoint in that they are readily adaptable and can be appropriately tuned for maximum fitness. Similarly, the increased mitochondrial copy number observed with some selections (e.g, ARC08-88; Figure 2C) could contribute to an enhanced stress response and high-grade resistance phenotype. The initial phenotypic response and resultant fading high-grade ATO/DSM1 resistance seen in ATO, MEN, and DSM1 selections can all be attributed to a combination of these genetic plasticity features, leading to metabolic plasticity with clear advantages for the parasite.

We next sought to estimate the worldwide frequency and distribution of mitochondrial heteroplasmy. We analyzed the publicly available *P. falciparum* data from the MalariaGEN Pf3k project (www.malariagen.net/pf3k)^28^ to uncover heteroplasmic mitochondrial diversity globally. After removing samples of multiple infections, lab lines, duplicates, and progenies of genetic crosses, we calculated the copy number of mitochondria number *Ĉ* of 1279 genomes using the sequencing depth differences between nuclear genome and mitochondrial genomes, with sampling-computation based correction of the sequencing coverage bias impacted by GC content^29^. For validation of our method, we estimated the copy number of nuclear chromosome as controls; and obtained the median copy number value of 1.00 (Figure 2B), consistent with the haploid genome of *P. falciparum* ^30^ In contrast, the median of mitochondrial copy number was 21.75 (Figure 2B). The majority of mitochondrial copy numbers from different samples ranged from 10 to 30, suggesting significant copy number variation in mitochondria globally (F-test p-value < 2.2e-16). The results are in agreement with recent reports of qPCR experiments^31^ which only used *cytochrome b* to estimate copy number.

To further quantify the level of heteroplasmy in parasite populations, a statistical measurement of maximum likelihood (PL score) of SNP detection based on VCF (Variant Call Format) output in the GATK pipeline^32^ (Figure 3A) was developed to detect allele frequency (AF). The lower AF value indicates lower levels of heteroplasmy, while AF = 1 indicates homoplasmy, either wild type or mutant. We scanned 1279 polymorphic isolates and observed heteroplasmy in the majority of polymorphic sites (n =1033, coverage > 200x) (Figure 3A & D). Our results showed that *P. falciparum* mt-diversity is currently underestimated by at least 3-fold (Poisson distributions, p < 0.001) without taking heteroplasmy into account. Geographically-specific polymorphisms exist in both homoplasmic and heteroplasmic SNPs (Figure 3B), suggesting important phenotypes might be associated with heteroplasmy. Interestingly, Cambodian *P. falciparum* isolates had a much higher rate of heteroplasmic SNPs than isolates from Ghana (Figure 3B). In addition, the mitochondrial copy number varied geographically with the highest copy numbers found in SE Asia where antimalarial drug resistance repeatedly emerges (Figure 3C). In agreement with widespread heteroplasmy in the global population, our study of deep-sequencing Asian parasite collections also revealed heteroplasmy, with Y268S as a low frequency common allele (Figure 2A & 3E).

**Figure 3.**
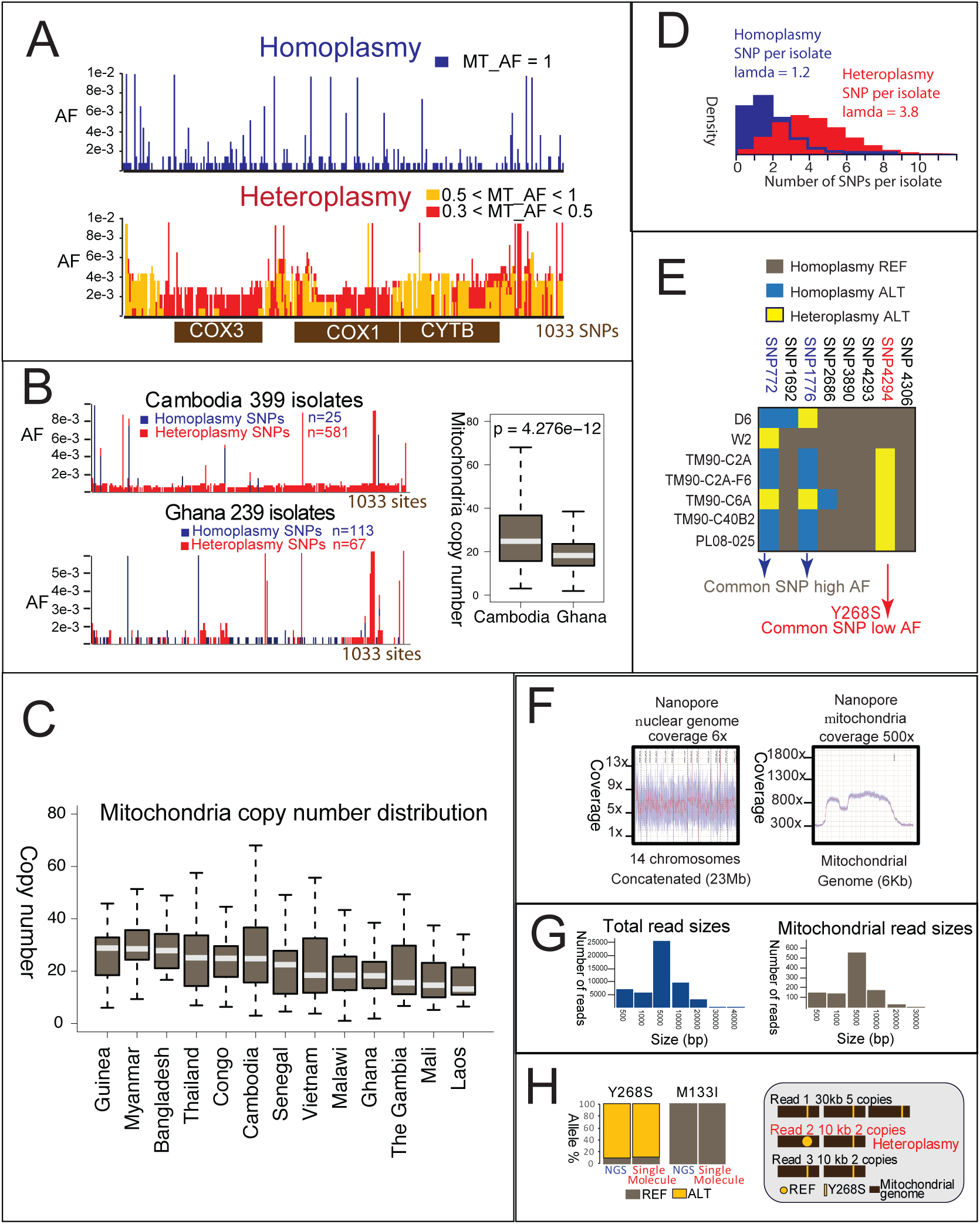
Analysis of the *P. falciparum* mitochondrial genetic diversity with data from the MalariaGEN Pf3K^28^ project and single molecule sequencing. A. Analysis of the heteroplasmic mitochondrial diversity in the 1279 parasite genome collection. A total of 1033 high-confidence SNPs with hundreds of mapped reads for support were studied. B. Geographical specific polymorphisms are found in both homoplasmy and heteroplasmy SNPs in Cambodia and Ghana. Cambodian isolates have higher diversity and higher mitochondrial copy numbers. C. Boxplot shows the copy number distribution of mitochondria in different regions. The regions are ranked based on the median value of copy number. The data from Nigeria is not shown here due to low sample number (n = 3). D. In the global population of 1279 parasites, number of SNPs per isolate is estimated with a Poisson distribution (Difference between homoplasmy and heteroplasmy pvalue < 0.001). E: Low level Y268S heteroplasmy is a common, low level SNP found in all patient admission isolates from the Phase II clinical trial, in contrast to high frequency, common heteroplasmic loci. F. Nanopore sequencing covers the nuclear genome at > 6x and mitochondrial genome at >500x. G Average reads length of Nanopore sequencing is 5 kb; and the longest read is 40 kb. H. Nanopore sequencing and NGS found the same allele frequencies. Y268S hetereplasmy is shown on a 20 kb single molecule reads with two tandemly arranged mitochondrial DNA units

Finally, to provide physical evidence of Y268S heteroplasmy, we performed single molecule sequencing with Nanopore^33^ on mtDNA of TM90-C2B. We obtained >500x coverage of mitochondrial genome and longest reads of >20 kb (Figures 3F and 3G). The single molecule sequencing resulted in the same levels of allele count with NGS sequencing with 90% Y268S in both systems (Figure 3H). Single reads consisting of both Y268S and references alleles were detected, thus supporting the heteroplasmy genetics of ATO resistance and mitochondrial diversity (Figure 3H).

Mitochondrial function in *P. falciparum* changes as the parasite transitions from asexual erythrocytic stages into gametocytes that are infectious to mosquitos. The increase in aerobic respiration is thought to block the transmission of cyt. *b* SNPs associated with ATO resistance^34^, although other studies demonstrated successful transmission of cyt. *b* M133I mutants^25^. Regardless, our data suggest heteroplasmy is transmitted and maintained as shown in the population diversity data, since both homoplasmic and heteroplasmic SNPs could be seen circulating in local populations (Figure 3B). Conceivably, heteroplasmy allows both neutral polymorphisms and alleles with fitness costs to be maintained at low frequencies and successfully transmitted. Additional fitness studies are required to further assess the transmission potential of the Y268S mutants with high copies of the mutant allele. The mechanism by which low-level diversity is maintained in organisms with similar mt-DNA structure/replication strategies has been studied extensively and is known as substoichiometric shifting. In plants, substoichiometric shifting produces unique subgenomic mitochondrial DNA molecules as the result of a recombination-based replication strategy, and can confer fitness advantages. It is unknown whether substoichiometric shifting can explain the maintenance of subgenomic mtDNA variants in *Plasmodium*, but it warrants further investigation.

In this study we provide multiple lines of evidence that mitochondrial heteroplasmy is the genetic mechanism underlying ATO resistance. These include *in vitro* drug selection, ATO resistance phenotype analysis, clinical isolate characterization, pyrosequencing analysis, mitochondrial copy number, and single molecule sequencing. Although current polymorphism studies represent a conservative estimate of heteroplasmy and mitochondrial copy numbers, our population analysis shows intriguing mitochondrial diversity exists in SE Asia, where resistance first emerged for many antimalarial drugs. Furthermore, increased mitochondrial copy number was observed in the same region, and also in parasites exposed to drug selection pressure *in vitro.* Our study revealed two novel aspects of malarial parasite mitochondrial genetics, i.e, heteroplasmy and copy number variations related to ATO resistance; these results will be crucial for developing other mitochondrial-targeting drugs, as well as combating drug resistance for eventual malaria elimination.

## Materials and Methods

### Parasite lines and cell culture

Admission and recrudescent parasite samples were collected from patients in the Phase II clinical trials upon admittance for treatment, and following failure of the treatment regimen (various dose regimens of atovaquone monotherapy or atovaquone/pyrimethamine combination therapy)^14^. The parasite history for paired admission and recrudescent isolates are outlined in Supplemental Table 1. Parasites were adapted to *in vitro* culturing and maintained according to the methods previously described by Trager and Jensen, with modifications first described by Webster *et al.*^35,36^ Parasites were maintained at 2% hematocrit in human O+ erythrocytes in RPMI 1640 (Invitrogen) medium containing 25 mM HEPES, 28 mM NaHCO_3_, 10% human type A positive plasma and incubated at 37°C in 5% O_2_, 5% CO_2_, and 90% N_2_ atmospheric conditions. Cultures were sustained with media changes three times per week and kept below 5% parasitemia with sub-culturing.

### Drugs and chemicals

Atovaquone (ATOV; 2-hydroxynapthoquinone), was purchased from Sigma (St. Louis, MO). Menoctone was synthesized and purified by the Manetsch laboratory at the University of South Florida, Department of Chemistry. DSM1 was kindly provided by Pradipsinh Rathod at the University of Washington. All compounds were used following dissolution in DMSO with final solvent concentrations less than 0.5%.

### Parasite EC_50_ determinations with hypoxanthine [^3^H] incorporation assay

The methods used were performed as described previously ^24^, with a modification of a 72-hour incubation period.

### Selection of atovaquone or menoctone resistant parasites *in vitro.*

In order to evaluate whether the genetic cryptic heteroplasmy background of parasite strains is essential to the development of the Y268S mutation conferring atovaquone resistance, we assessed the resistance potential of admission isolate clones TM90-C2A-F6, TM90-C40B2, and ARC08-88-8A. TM90-C2A and TM90-C40 were taken from patients prior to treatment and later recrudesced with Y268S mutations in cyt. *b* following atovaquone monotherapy regimens (Supplementary Table 1). TM90-C2A, TM90-C40, and ARC08-88 were sub-cloned by limiting dilution^37^ prior to any drug selections, and sub-clones TM90-C2A-F6, TM90-C40B2, and ARC08-88-8A were used for all drug selections. Sub-cloning the parasites prior to drug selections was necessary in order to provide an isogenic background as well as to maximize phenotypic stability, as many of these parasites experienced more widely fluctuating EC_50_ values to mitochondrial inhibitors (4-8 fold) than control parasites (> 3 fold). We hypothesize the phenotypic fluctuations are a function of the genotypic plasticity within the population as a whole, where successive replication rounds vary somewhat in Y268S frequency. ARC08-88 was originally obtained from the World Health Organization Global Plan Artemisinin Resistance Containment consortium, and was used to demonstrate the development of atovaquone resistance from a parasite outside the Phase II studies of atovaquone in Thailand that had cryptic Y268S heteroplasmy. TM90-C2A, TM90-C40B2, and ARC08-88-8A were grown from earliest available cryopreserves to 10^8^ and seeded into 25 ml flasks in triplicate. The complete medium contained approximately 10x EC50 atovaquone (10 nM) or 10x EC50 menoctone (1.5 μM) with media changed twice per week, and split 1:2 with fresh erythrocytes every 10 days to maintain 2% hematocrit. Parasites were considered “recovered” from drug selection when parasite densities reached 2% parasitemia and sustained growth under continuous drug pressure. All parasites had the cytochrome *b* gene sequenced to look for possible mutations developed during drug pressure.

### Parasite genomic DNA isolation

*P. falciparum-infected* erythrocytes were treated with 0.05% saponin for 10 min and genomic DNA (gDNA) was extracted with the Qiagen DNeasy Kit according to manufacturer’s protocols. Unless described otherwise, gDNA was harvested from parasites from earliest possible cryopreservation dates to best preserve phenotypes and genotypes.

### PCR and cytochrome *b* and DHODH gene sequencing

All *P. falciparum* cytochrome *b* PCR products were amplified using primers cytbFOR 5’—TGCCTAGACGTATTCCTG—3’ and cytbREV 5’—GAAGCATCCATCTACAGC—3’. PCRs were amplified using Phusion HS II High-Fidelity PCR Master Mix (ThermoFisher Scientific) with ∼20 ng parasite gDNA template, according to manufacturer’s instructions with the following program: 98°C—30s initial denaturation step, then 35 cycles: (98°C—10s, 54°C—40s, 72°C—30s) and a final extension of 72°C for 7 min. PCR products were confirmed as a single, discrete band of 1,382 bp length on a 1% agarose gel then subsequently purified using the Qiagen PCR Purification Kit according to manufacturer’s instructions. Purified PCR products were prepared for Sanger sequencing service at Genewiz (Genewiz, South Plainfield, NJ) using the following sequencing primers: pf-cytb-SEQFOR1: 5’—GTGGAGGATATACTGTGAGTG—3’, pf-cytb-SEQFOR2: 5’— TACAGCTCCCAAGCAAAC—3’, pf-cytb-SEQREV1: 5’—GACATAACCAACGAAAGCAG—3’, and pf-cytb-SEQREV2: 5’—GTTCCGCTCAATACTCAG—3’. The *dhodh* gene was sequenced similarly using primers described in Ross *et al^19^*. Sample sequences were analyzed and aligned using ApE (A Plasmid Editor) software, and mapped to the Pf-3D7 *cytochrome b* gene annotated on Plasmodb.org for mutation detection.

#### Sequencing to identify SNPs in mitochondrial genes

We sequenced multiple candidate genes in the mtETC to determine if additional SNPs were associated with the drug resistance spectrum phenotypes. All PCR reactions were set up similarly to the sequencing of the cytochrome *b* gene above. Candidate genes included PF3D7_0915000 (NDH2), PF3D7_0603300 (DHODH), PF3D7_0523100 (Core 1), PF3D7_093360 (Core 2), PF3D7_1462700 (cyt. *c_1_*), PF3D7_1439400 (Rieske), PF3D7_1426900 (QCR6), PF3D7_1012300 (QCR7), and the three mt-encoded genes: MAL_MITO_3 (cyt. b), MAL_MITO_1 (*coxlll)*, MAL_MITO_2 (*coxl).* For the primers in Extended Data Table 5, those labeled PCR FOR and PCR REV were used in PCR amplifications, and SEQ PR denotes primers used to sequence the amplification in its entirety.

### Pyrosequencing of Y268S allele

The Pyromark Q96 ID system was used for the detection of single-nucleotide polymorphism (SNP) for Y268S detection in Pf-cytochrome b, with Qiagen Pyromark Gold Q96 reagents and buffers along with streptavidin sepharose beads (GE Healthcare). All template and reaction components were prepared according to manufacturer’s protocols. Pyrosequencing primers were designed using Pyromark Assay Design Software. Primers for the initial PCR reaction were amplified with PFcytb_pyro_Biotin_FOR 5’—Biotin-ACCATGGGGTCAAATGAGTTAT—3’ and PFcytb_pyro_REV 5’—AGCTGGTTTACTTGGAACAGTTTT—3’ as 50 μL reactions with 25 μL 2X Phusion Hot Start II HF PCR Master Mix, 0.2 μM primer concentrations, ∼10-50 ng template gDNA, brought to 50 μL total volume with nuclease-free water, with the following thermocycling conditions: initial denaturation of 98°C for 30s, 55 cycles of 98°C for 30s, 53°C for 5s, and 72°C for 8s. All parasites resulting from *in vitro* drug selections had gDNA harvested immediately following parasite recovery to 2% parasitemia, which provided insight into the early period of resistance where more extreme phenotypes were observed in EC_50_. Subsequent PCRs were run on 1.5% agarose gels to confirm a single discrete band without excess primer present, as unconsumed primer has been shown to interact with pyrosequencing primers to contribute to a background signal in no template controls, and was minimized by using low primer concentrations and using a high cycle number to exhaust primers. The Pyromark pyrosequencing assay was performed according to manufacturer’s protocols with pyrosequencing primer PFcytb_seq_assay_REV 5’— TGGAACAGTTTTTAACATTG—3’. Each parasite gDNA sample was initially amplified independently in triplicate, and had two technical replicates per reaction (25 μL PCR per pyrosequencing reaction) on the Pyromark Q96 ID for a total of at least 6 pyrosequencing runs per parasite gDNA template. Allele frequencies were analyzed by Pyromark ID software in allele quantification mode.

#### Pyrosequencing Y268S assay standard curve

Since all Y268S mutant genotypes still contained some small quantities of wild-type allele, we chose to use the parasite with the highest percentage Y268S mutant, TM90-C50B5 gDNA (99.52% mutant) and D6 (0% mutant) were mixed at 10% increments from 0% wild type gDNA + 100% wild type gDNA, adding in additional increment mixtures at the lower 5% and upper 95%, with 1% increments to look at the sensitivity of detection. These ratio wild type:mutant gDNA mixtures were made independently three times and then used in subsequent PCR reactions and pyrosequencing reactions to generate the standard curve in Figure 1A.

#### DHODH Copy Number Variation Quantitative PCR (qPCR)

Pf-DHODH copy number was determined using the DHODH qPCR primers previously described by Guler *et al*^27^ and the LDH-T1 FOR/REV control primers from Chavchich *et al*.^38^ using Brilliant II/III SYBR Green Master Mix with ROX and the Mx3005P qPCR machine (Applied Biosystems). The relative copy number of DHODH was determined for 0.1 ng of gDNA and normalized to the LDH gene using the ΔΔC_T_ method^39^.

#### Deep sequencing analysis and data mining

For deep sequencing of clinical and *in vitro* selected parasite genomes, we ran samples on the Illumina HiSeq 3000 with the 150 cycle protocol (PCR-free) and used >1 ug purified DNA as starting material, and reached 10,000-30,000 x coverage of mtDNA. Each allele must have a minimum of 100 high quality reads (QC score > 60) mapped to the loci to be identified as a rare allele. As controls, we examined all loci with the same stringency, and our method did not recover any low frequency heteroplasmy in the vast majority of the loci (n=5954), other than a few loci including Y268S.

For computational mining of the sites with heteroplasmy in the global parasite populations, we developed a maximum likelihood based method to differentiate high confidence heteroplasmic sites from background error count. We use a statistical measurement of maximum likelihood (PL score) of SNP detection based on VCF (variant call format) output in GATK pipeline^32^ to detect allele frequency (AF). First, we obtained the genotype likelihood (GL) for the given loci, with GL defined as LOG10 scaled likelihoods for all possible genotypes given the set of alleles defined in the REF and ALT fields, as specified by GATK protocols^40,41^. Then, the phred-scaled genotype likelihood (PL) score is obtained. Only the PL score of being heteroplasmic vs being homoplasmic, at a given locus, larger than 6 (i.e, Pvalue < 0.000001), is considered a true heteroplasmic site. The lower AF value indicates lower level of heteroplasmy, while AF = 1 indicates homoplasmy, either wild type or mutants.

To independently confirm our computational mining methods, we examined 100 isolates from the population data sets. We retrieved the raw data mitochondrial BAM files, and examined that the heteroplasmic sites are correctly identified by the maximum likelihood method, as shown by more than a hundred high quality reads in the mapping alignments.

We used the sequencing depth differences to estimate the copy number of query chromosome from sequencing signal. Here, we assume the PCR amplification level is same for each chromosome copy. It has been reported the copy number of nuclear chromosome is 1 during asexual stage in *Plasmodium^30^.* Therefore, the sequencing depth difference between input and reference nuclear chromosome is the estimated copy number of query chromosome (equation 1).

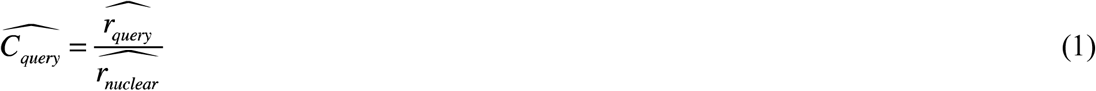

where 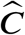 is the estimated copy number and 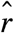 is the estimated sequencing signal. To find the best estimation of sequencing signal, we assume that the true sequencing reads signal of a whole specific chromosome follows a uniform distribution. Then we calculated the optimal sequencing depth with

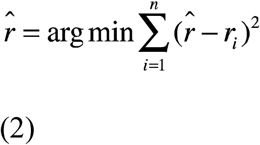

Here, 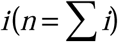 is a given segment on the chromosome; and the sequencing signal *r_i_* is calculated as

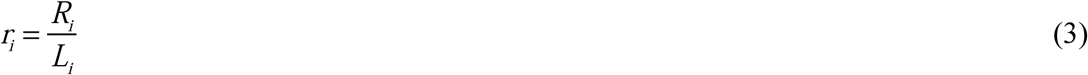

where *R_i_* is the total reads mapped on segment *i, L_i_* is the segment length. The final estimation 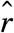 from equation 2 is

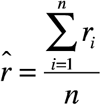

### Nanopore mtDNA sequencing

#### Isolation of mitochondrial DNA in TM90-C2B

Parasites were grown to 10% parasitemia in 50 μL culture volume, then synchronized with 5% sorbitol twice, spaced 4 hours apart, followed by MACS column purification to remove any residual trophozoites and schizonts so that only ring stages were collected to obtain non-replicative mtDNA molecule forms only. Red blood cells were then lysed using 0.1% saponin in 1x PBS, washed 3x in PBS, and mitochondrial DNA was isolated from parasite material using the Abcam Mitochondrial DNA Isolation Kit using the manufacturer’s instructions, with the exception of using needle passage to homogenize cells. Isolated mitochondrial DNA was used for subsequent Nanopore library preparation and sequencing.

#### Library Preparation and Sequencing

1 μg of parasite mitochondrial DNA (RNAse treated) was subsequently treated with protease and mtDNA eluted in 45 μL nuclease-free water (NFW). End-repair and dA-tail of mtDNA was performed by adding 7 μL Ultra II End-Prep buffer, 3 μL Ultra II End-Prep enzyme mix (NEBNext Ultra II End-Repair/dA-tailing Module, New England BioLabs), and 5 μL NFW. The mix was incubated for 5 minutes at 20 °C and 5 minutes at 65 °C using a thermocycler. The end-prep reaction cleanup was performed by adding 60 μL of resuspended AMPure XP beads and mtDNA was eluted in 31 μL NFW. A 1 μL aliquot from the elute was quantified by fluorometry (Qubit) to ensure ≥700 ng end-prepped mtDNA was retained.

Adapter ligation was performed by adding 10 μL of Adapter Mix, 2 μL HP Adapter (SQK-NSK007 Nanopore sequencing Kit, Oxford Nanopore Technologies), 50 μL NEB Blunt/TA Master Mix (NEB, cat no M0367), and 8 μL NFW to 30 μL dA-tailed mtDNA, mixing gently and incubating at room temperature for 10 minutes. 1 μL of HP Tether (SQK-NSK007) was added to the mix and incubated for 10 minutes (RT). Library purification of the adapted and tethered mtDNA was performed by adding MyOne C1 Streptavidin Beads, incubated for 5 minutes at room temperature and resuspended the pellet in 150 μL Bead Binding Buffer (SQK-NSK007). The purified-ligated mtDNA pellet was resuspended in 25 μL Elution Buffer (SQK-NSK007), incubated for 10 minutes at 37 °C and the 25 μL elute (Pre-sequencing Mix) was transferred into a new tube. A 1 μL aliquot was quantified by fluorometry (Qubit) to ensure ≥ 500 ng of adapted and tethered mtDNA was retained.

The MinION Flow Cell (R9) was primed twice prior to loading sample with 500 μL flow cell priming mix made of 1:1 ratio of Running Buffer with Fuel Mix 1 (RBF1) and NFW. The library for loading was prepared by adding 12 μL adapted and tethered library, 75 μL RBF1 and 63 μL NFW. Using a P-1000 tip set to 150 μl, the library was loaded into the flow cell keeping the pipette vertical. The flow cell was run on the 48ᛱh 2D protocol. Data Analysis

Nanopore sequencing reads were processed using the Metrichor cloud platform 2D workflow. Only the reads (Fast5 files) that passed Metrichor quality cutoffs were converted into fastq format using poretools (v0.5)^42^. The reads were aligned against the Plasmodium falciparum 3D7 (PlasmoDB v28)^43^ reference genome using the BWA-MEM (v0.7.15)^44^ with settings “-x ont2d”. The aligned data was used to generate a.bam file using SAMtools (v1.3.1)^45^ and BCFtools (v1.3.1). Finally, alignments were visually inspected using IGV viewer^46^ in an attempt to trace back the % Y268S mutation in the parasite.

---

**Extended Data Table 1.** Cytochrome b mutations induced by drug selection with atovaquone (ATO) and menoctone (MEN) in vitro

| Parasite | Drug | Cyt. *b* mutation(s) | Source |
| --- | --- | --- | --- |
| D6 | ATO | No resistance | Rathod *et al.* 1997 |
| 3D7 |  | M133I<br>M133I & P275T<br>M133I & K272R<br>M133I & G280D<br>L283I & V284K | Korsinczky *et al.* 2000 |
| 3D7 |  | M133V<br>M133I<br>M133I & L144S<br>F267V | Bopp *et al.* 2013 |
| K1 |  | M133I<br>M133I & G280D | Schwobel *et al.* 2003 |
| AT200 |  | M133I<br>M133I & L271F | Schwobel *et al.* 2003 |
| TM90-C2A |  | Y268S | This study |
| TM90-C40B2 |  | Y268S | This study |
| ARC08-88-8A |  | Y268S | This study |
| W2 | MEN | M133I | This study |
| TM90-C2A |  | Y268S | This study |

**Extended Data Table 2.**
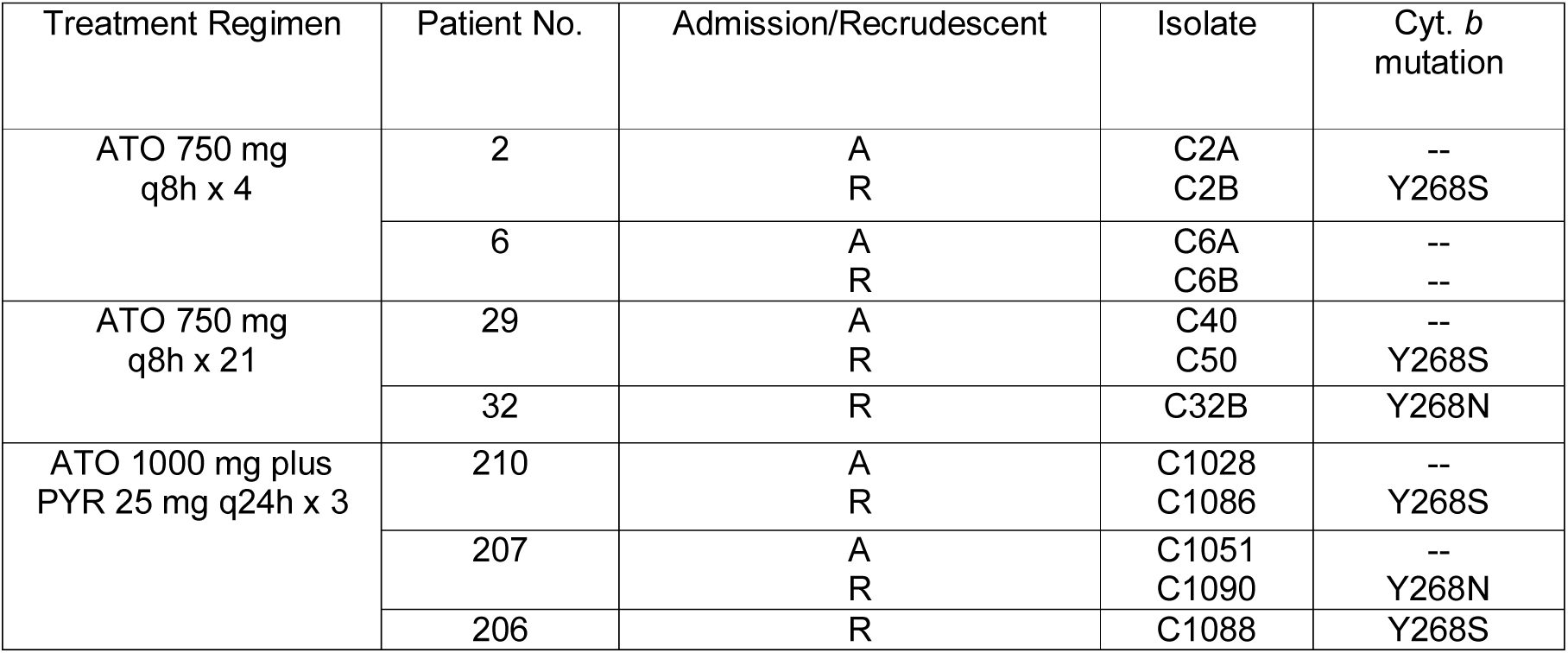
Treatment regimens and cytochrome b (cyt b) genotypes of admission (A) and recrudescent (R) isolates of *Plasmodium falciparum* from Phase II clinical studies with atovaquone (ATO) alone or in combination with pyrimethamine (PYR)^14^

| Treatment Regimen | Patient No. | Admission/Recrudescence | Isolate | Cyt. b mutation |
| --- | --- | --- | --- | --- |
| ATO 750 mg<br>q8h x 4 | 2 | A<br>R | C2A<br>C2B | --<br>Y268S |
|  | 6 | A<br>R | C6A<br>C6B | --<br>-- |
| ATO 750 mg<br>q8h x 21 | 29 | A<br>R | C40<br>C50 | --<br>Y268S |
|  | 32 | R | C32B | Y268N |
| ATO 1000 mg plus<br>PYR 25 mg q24h x 3 | 210 | A<br>R | C1028<br>C1086 | --<br>Y268S |
|  | 207 | A<br>R | C1051<br>C1090 | --<br>Y268N |
|  | 206 | R | C1088 | Y268S |

**Extended Data Table 3.** Sanger sequence results of candidate mtETC resistance genes in patient isolates and reference clones of *P. falciparum*

| Gene ID | Parasite names (abbreviated) |  |  |  |  |  |  |  |  |  |  |  |
| --- | --- | --- | --- | --- | --- | --- | --- | --- | --- | --- | --- | --- |
|  | W2 | D6 | C2A | C2B | C6A | C6B | C40B2 | C50B5 | C1051 | C1090 | C1086 | C1088 |
| PF3D7_0915000<br>(NDH2) | -- | -- | -- | -- | -- | -- | -- | -- | ND | ND | -- | -- |
| PF3D7_0603300<br>(DHODH) | -- | -- | -- | -- | -- | -- | -- | -- | -- | -- | -- | -- |
| MAL_MITO_1<br>(COXIII, Complex IV) | -- | I239V | I239V | I239V | -- | -- | I239V | I239V | I239V | -- | -- | -- |
| MAL_MITO_2<br>(COXI, Complex IV) | -- | -- | -- | -- | -- | -- | -- | -- | -- | -- | -- | -- |
| MAL_MITO_3<br>(Cyt. *b*) | -- | -- | -- | Y268S | -- | -- | Y268S | Y268S | -- | Y268N | Y268S | Y268S |
| PF3D7_0523100<br>(Core 1, Complex III) | -- | -- | -- | -- | -- | -- | -- | -- | ND | ND | -- | -- |
| PF3D7_093360<br>(Core 2, Complex III) | -- | -- | -- | -- | -- | -- | -- | -- | ND | ND | -- | -- |
| PF3D7_1439400<br>(Rieske, Complex III) | -- | -- | -- | -- | -- | -- | -- | -- | ND | ND | -- | -- |
| PF3D7_1426900<br>(QCR6, Complex III) | -- | -- | -- | -- | -- | -- | -- | -- | ND | ND | -- | -- |
| PF3D7_1012300<br>(QCR7, Complex III) | -- | -- | -- | -- | -- | -- | -- | -- | ND | ND | -- | -- |
| PF3D7_1462700<br>(Cyt. *c*<sub>1</sub>) | -- | -- | -- | -- | -- | -- | -- | -- | ND | ND | ND | -- |

-- Indicates wild type sequence; ND – not determined

**Extended Data Table 4.** Genotypes of DSM1 drug selections immediately following recovery

| Genotypes of DSM1 drug selections immediately following recovery |  |  |  |  |
| --- | --- | --- | --- | --- |
| DSM1-selected populations | *dhodh* copy number | PFF_0160C (*dhodh*) | MALMITO_3 (cyt. *b*) | MALMITO_1 (*coxIII*) |
| C2A+10xDSM1-1A | 3 | WT | WT | I239V |
| C2A+10x DSM1-2B | 1 | R265G | Y268S/WT | I239V |
| C2B+2xDSM-1A | 2 | WT | Y268S/WT | I239V |
| C2B+2xDSM1-1B | 2 | WT | Y268S/WT | I239V |
| C2B+2xDSM1-3B | 1.5 | WT | Y268S/WT | I239V |
| C2B+2xDSM1-2C | 1 | WT | Y268S/WT | I239V |

**Extended Data Table 5.**
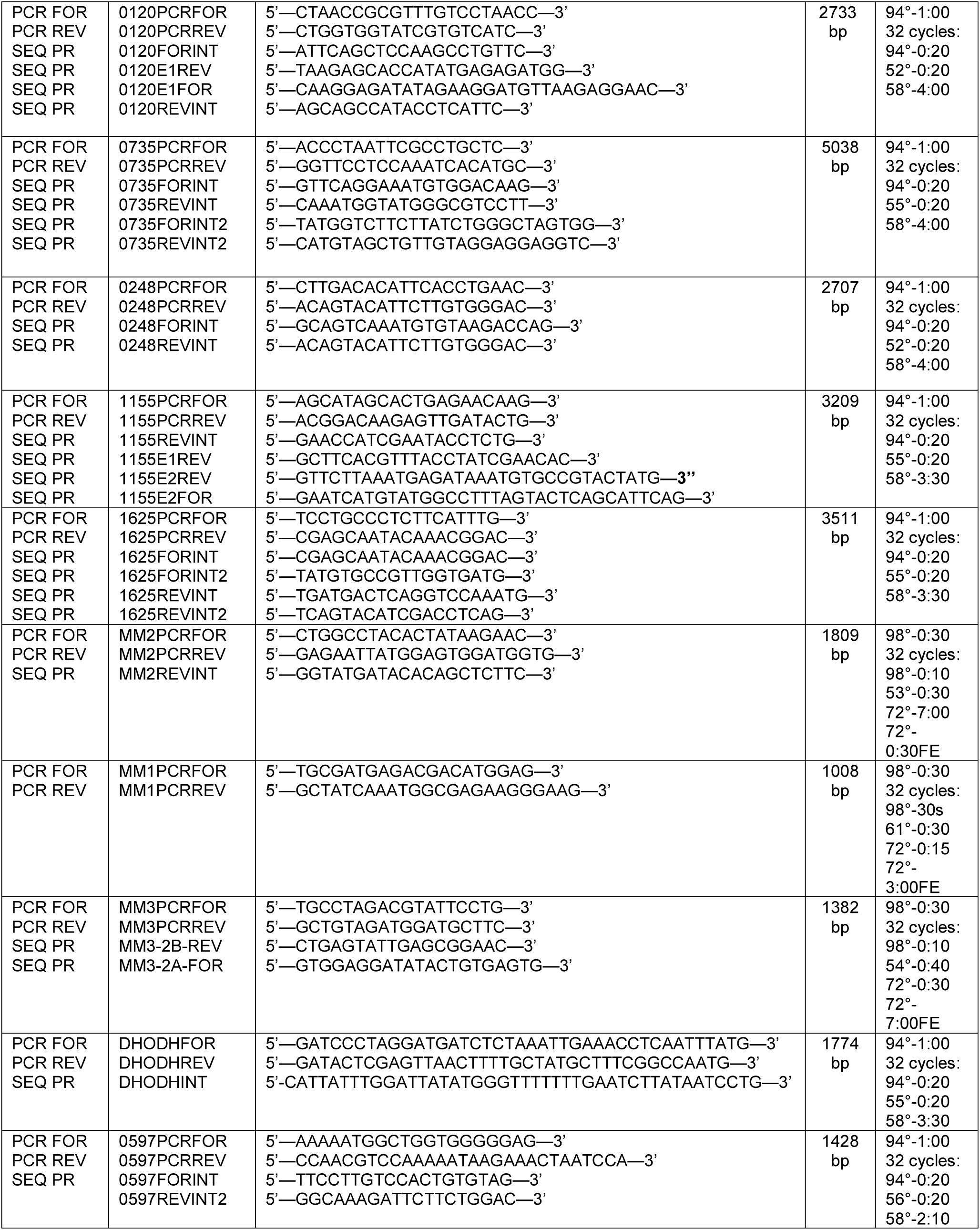
PCR Primers and Programs Used inmtETC Sequencing

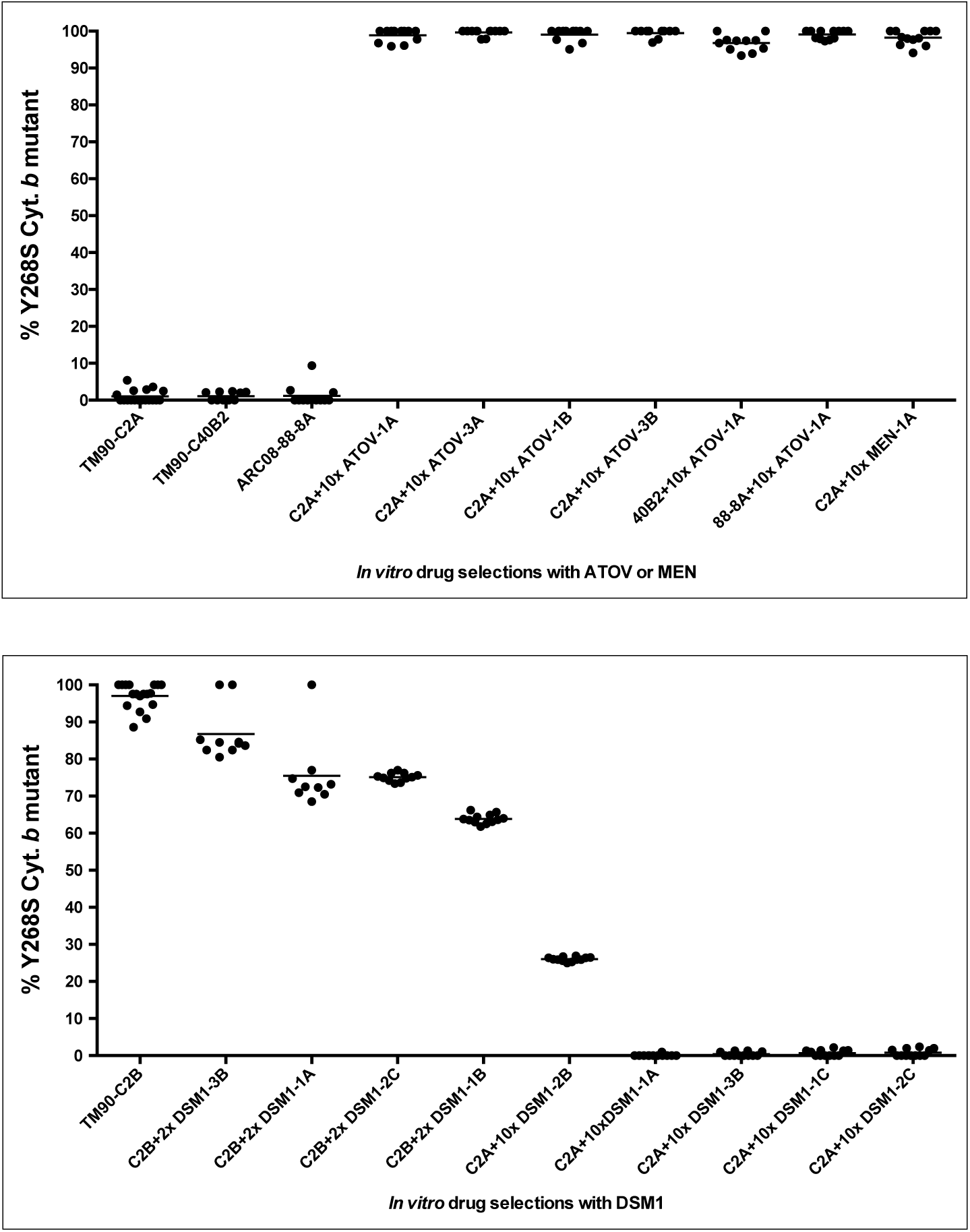
Extended Data Figure 1.

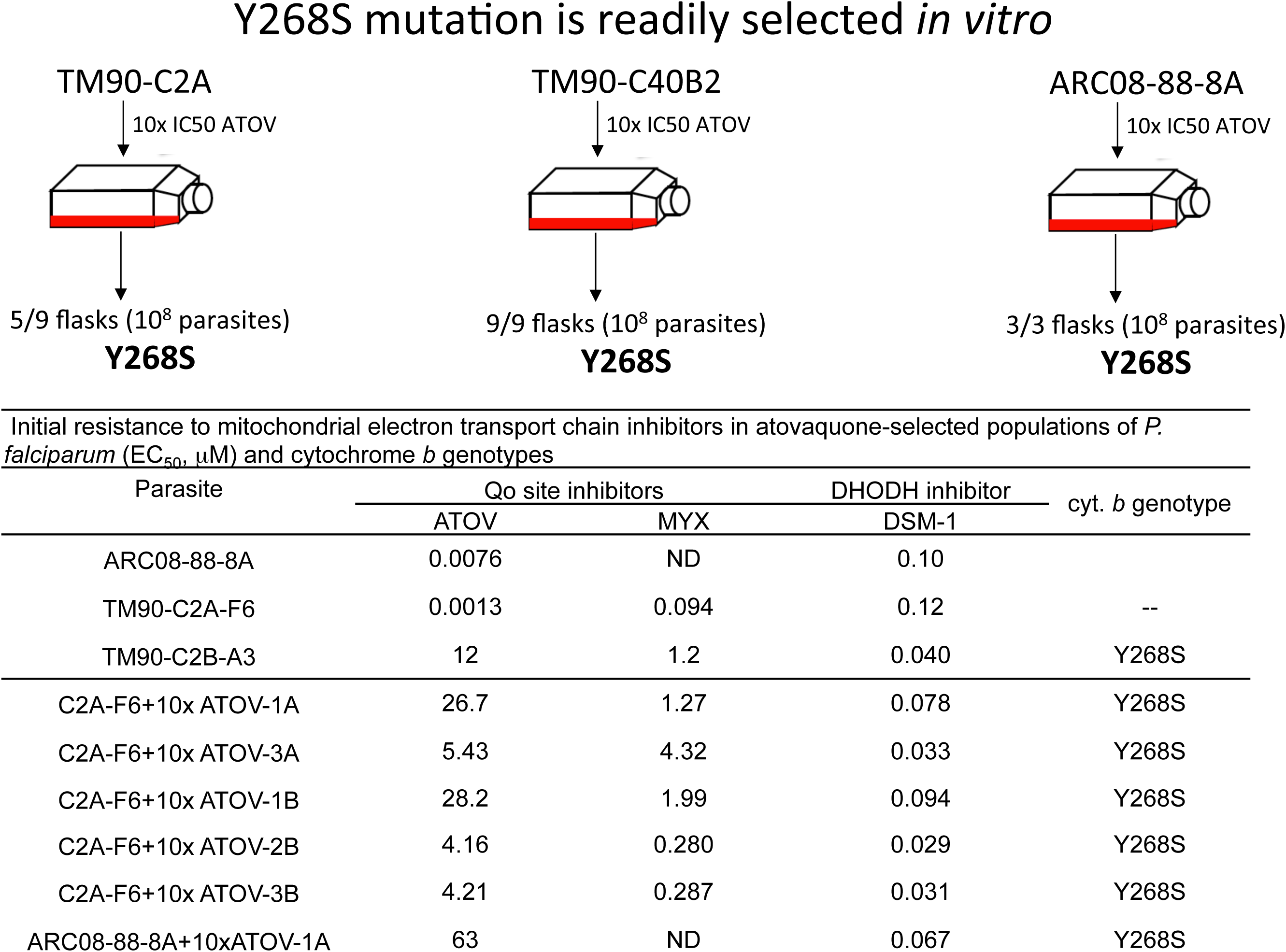
Extended Data Figure 2.

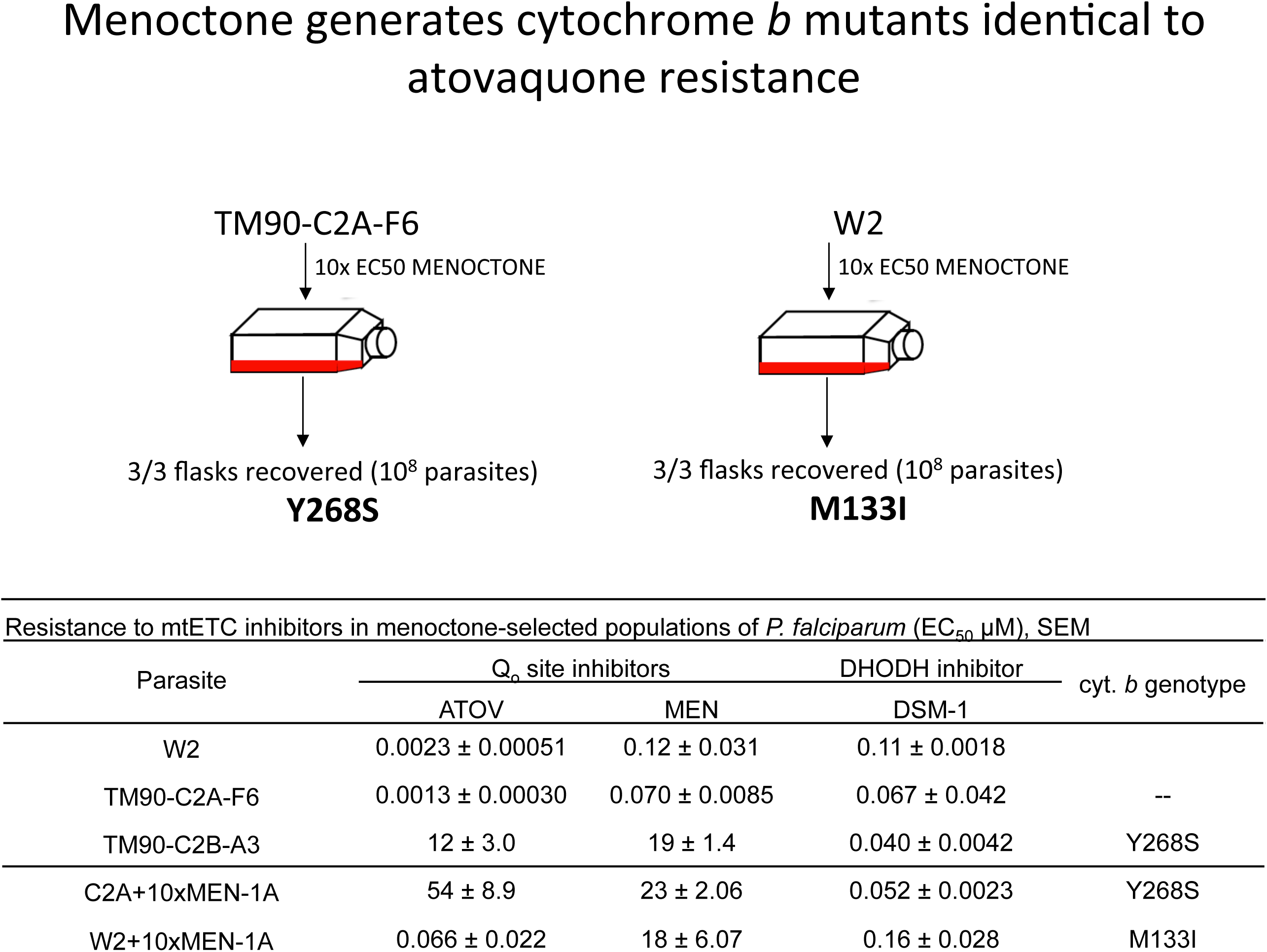
Extended Data Figure 3.

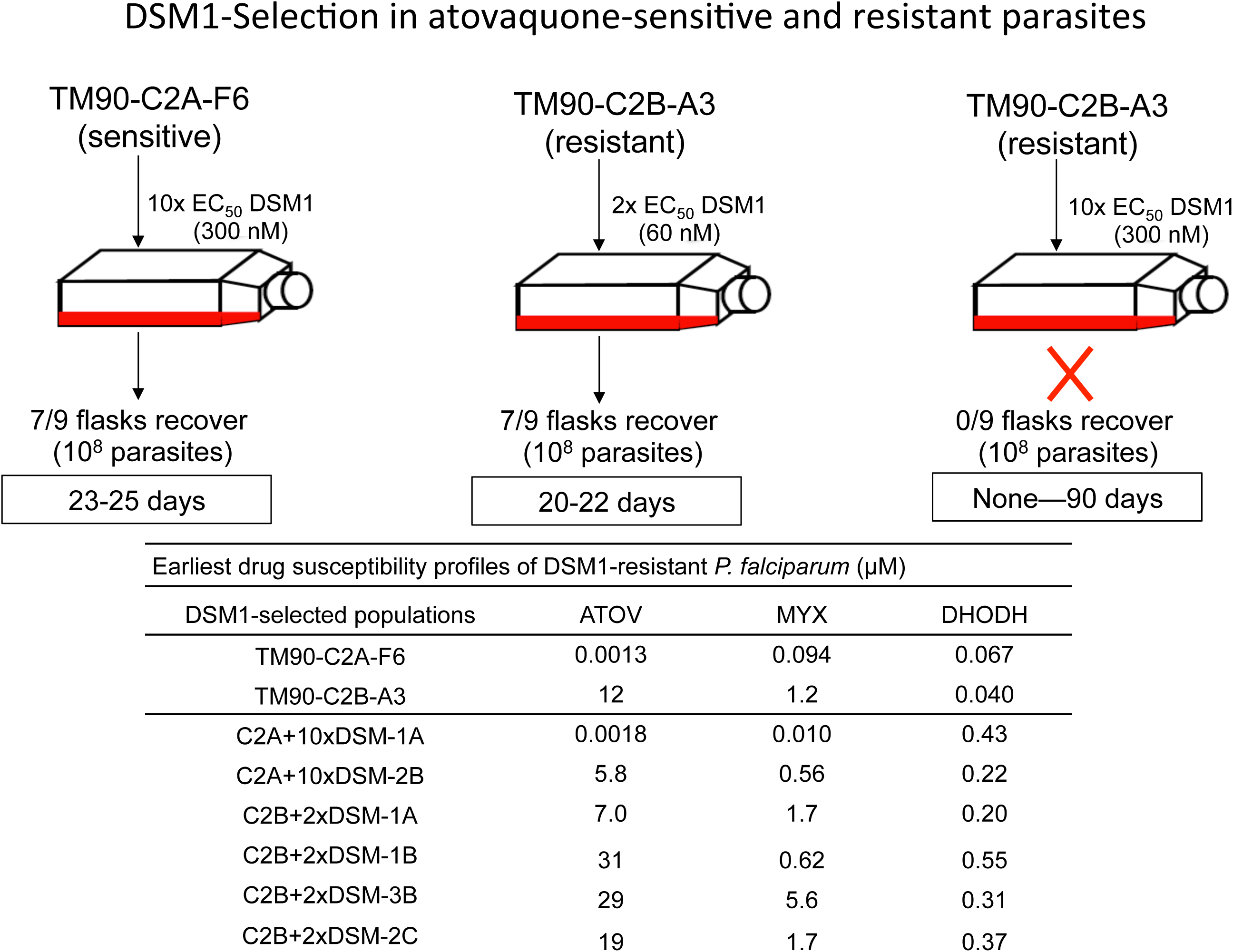
Extended Data Figure 4.

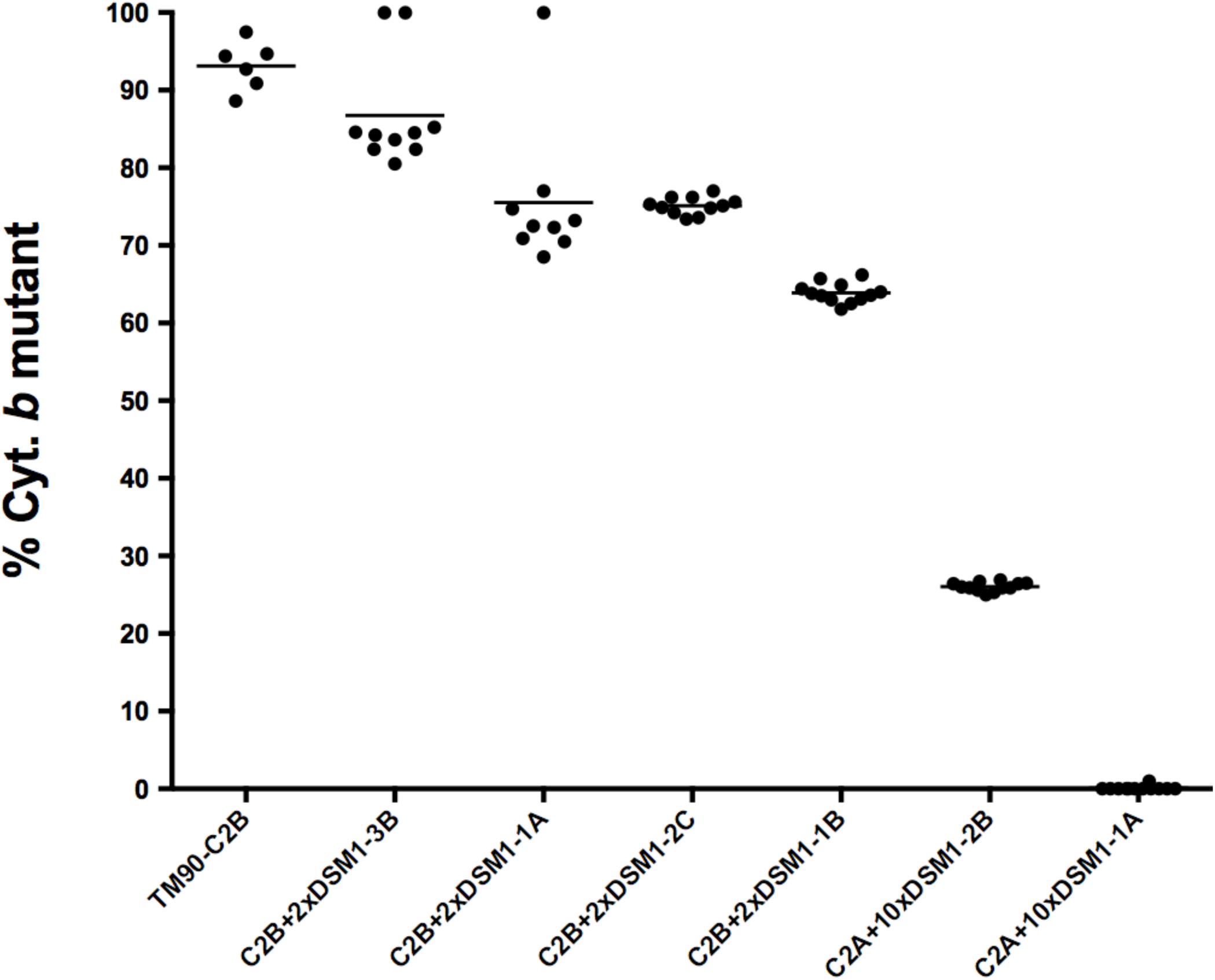
Extended Data Figure 5.

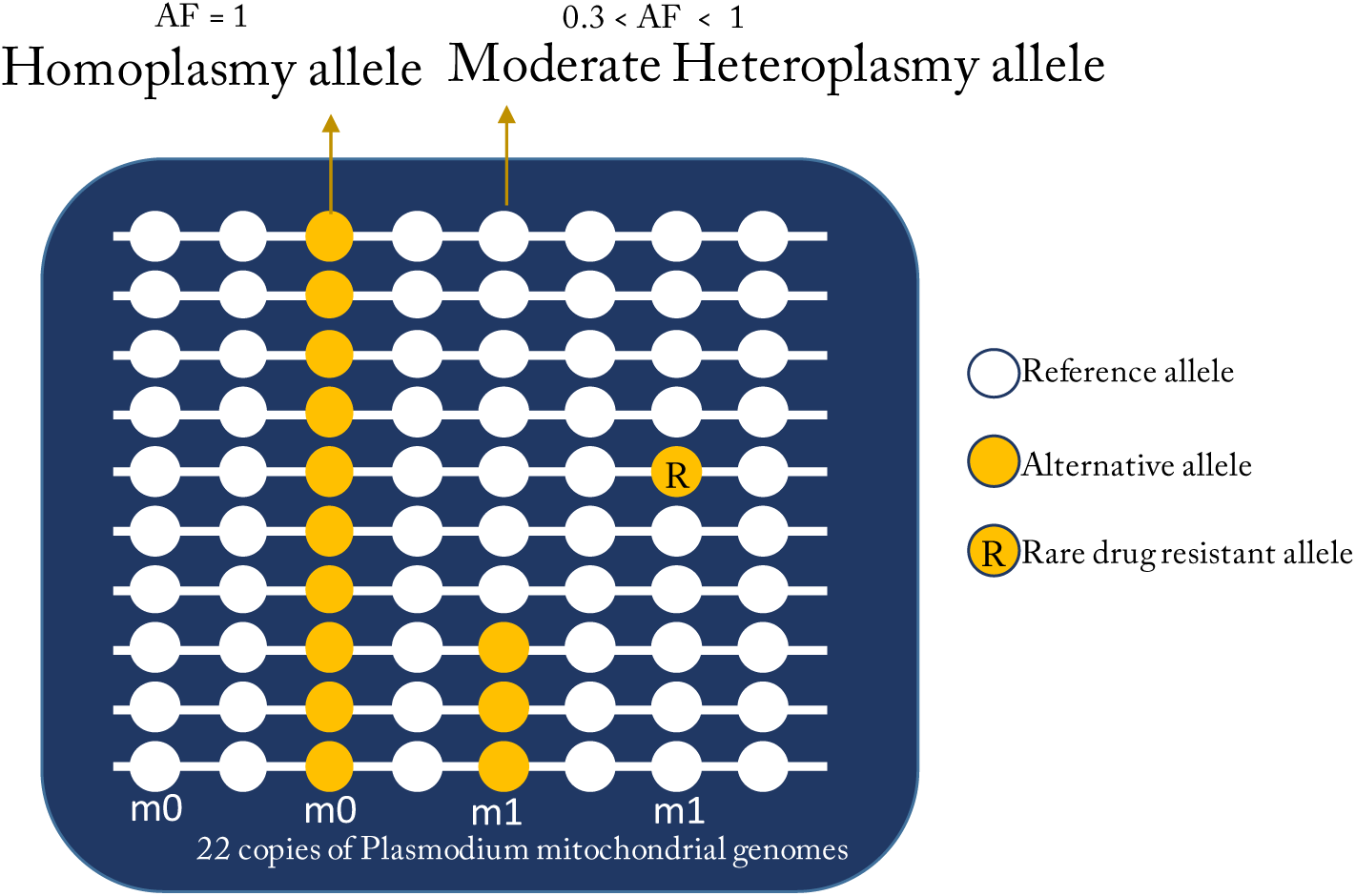
Extended Data Figure 6.

